# Development of Narrow Spectrum ATP-competitive Kinase Inhibitors as Probes for BIKE and AAK1

**DOI:** 10.1101/094631

**Authors:** Rafael M. Couñago, Alison D. Axtman, Stephen J. Capuzzi, Hátylas Azevedo, David H. Drewry, Jonathan M. Elkins, Opher Gileadi, Cristiano R. W. Guimarães, Alessandra Mascarello, Ricardo A. M. Serafim, Carrow I. Wells, Timothy M. Willson, William J. Zuercher

**Affiliations:** Structural Genomics Consortium, Universidade Estadual de Campinas – UNICAMP, Campinas, SP, Brazil.; Structural Genomics Consortium, UNC Eshelman School of Pharmacy, University of North Carolina at Chapel Hill, Chapel Hill, NC, USA.; Division of Chemical Biology and Medicinal Chemistry, UNC Eshelman School of Pharmacy, University of North Carolina at Chapel Hill, Chapel Hill, NC, USA.; Aché Laboratórios Farmacêuticos SA, Guarulhos, SP, Brazil.; Structural Genomics Consortium and Target Discovery Institute, Nuffield Department of Clinical Medicine, University of Oxford, Old Road Campus Research Building, Oxford, OX3 7DQ, UK.

## Abstract

Understanding the structural determinants of inhibitor selectivity would facilitate the design and preparation of kinase probes. We describe a pair of matched compounds differing only by one degree of saturation but showing dramatic differential activities at select kinases. We utilized x-ray crystallography and computational analysis to rationalize the basis of the differential activity.

## INTRODUCTION

The human protein kinases are a family of over 500 enzymes that are involved in critical aspects of cell signaling but are often dysregulated in disease, especially cancer [1]. Protein kinases catalyze the transfer of phosphate from the cofactor ATP to their substrate proteins. Although ATP-competitive protein kinase inhibitors have been successfully developed to treat a range of cancers, inflammatory and fibrotic diseases, the function of a large number of the enzymes remains unknown [2]. Potent and selective small molecule inhibitors are powerful tools to probe the biology of these understudied kinases [3].

Because ATP-competitive inhibitors bind to a common active site, they often show activity across multiple kinases. For example, staurosporine inhibits more than 200 protein kinases. These broad spectrum inhibitors are useful for initial biochemical characterization but have little utility as probes of kinase biology [4]. By contrast, other ATP-competitive inhibitors possess activity on only a handful of kinases. These narrow spectrum inhibitors have found utility in exploring the function of several historically understudied kinases [5].

To strengthen the conclusions drawn from use of narrow spectrum kinase inhibitors, a recently established best practice is to use the probe molecule in parallel with a negative control, a closely-related analogue that is inactive for the target kinase [6]. Demonstration of divergent results between the probe inhibitor and its negative control increases confidence that the engagement of the target kinase was responsible for the observed biology.

For ATP-competitive kinase inhibitors, a systematic approach to develop negative control analogs is to synthesize a molecule that lacks the ability to form the critical hinge binding interaction, typically composed of one or more hydrogen bonds with the highly conserved hinge region in the enzyme [7]. These modifications result in “kinase dead” analogues. Unfortunately, kinase dead analogues often lose affinity not only for the primary target kinase but also for all other off-target kinases. New strategies to create kinase probe negative controls that lose affinity selectively for their target kinase are needed.

Here we describe a pair of compounds in which a minor structural change—namely, the inclusion of a cyclopropyl versus an isopropyl substituent proximal to the hinge-binding moiety—leads to unexpectedly large changes in activity within a small subset of protein kinases but no change against many others. We used X-ray crystallography and computational models to study the molecular basis for these differential effects on affinity within the target and off-target kinases in an attempt to define general approaches to design selective negative control analogs.

## RESULTS

As part of a large-scale selectivity screen of ATP-mimetic kinase inhibitors, we identified a pair of very closely related 3-acylaminoindazoles (**1** and **2**) that displayed a narrow binding spectrum in the KINOMEscan panel of over 400 human protein kinase assays at an inhibitor concentration of 1 μM (**S1 Table**). Structurally, **1** and **2** differed only by the addition of two hydrogen atoms in the change from a cyclopropyl to isopropyl carboxamide. **1** and **2** had several common kinase targets but also showed some degree of divergence in kinase affinity profiles. A more detailed analysis of their binding characteristics revealed that **1** had submicromolar affinity for 11 kinases and weaker affinity for 2 additional kinases (**Table 1**), while **2** showed submicromolar affinity for 7/11 of the primary kinase targets and 1/2 of the minor targets. For most of these kinases, only marginal differences in potency were observed between the analogs (<3-fold). However, four kinases (AAK1, BIKE, RIOK1, and RIOK3) displayed remarkable differential affinity between the cyclopropyl analog **1** and isopropyl analog **2** with a preference for the former (27–150-fold).

**Table 1.**
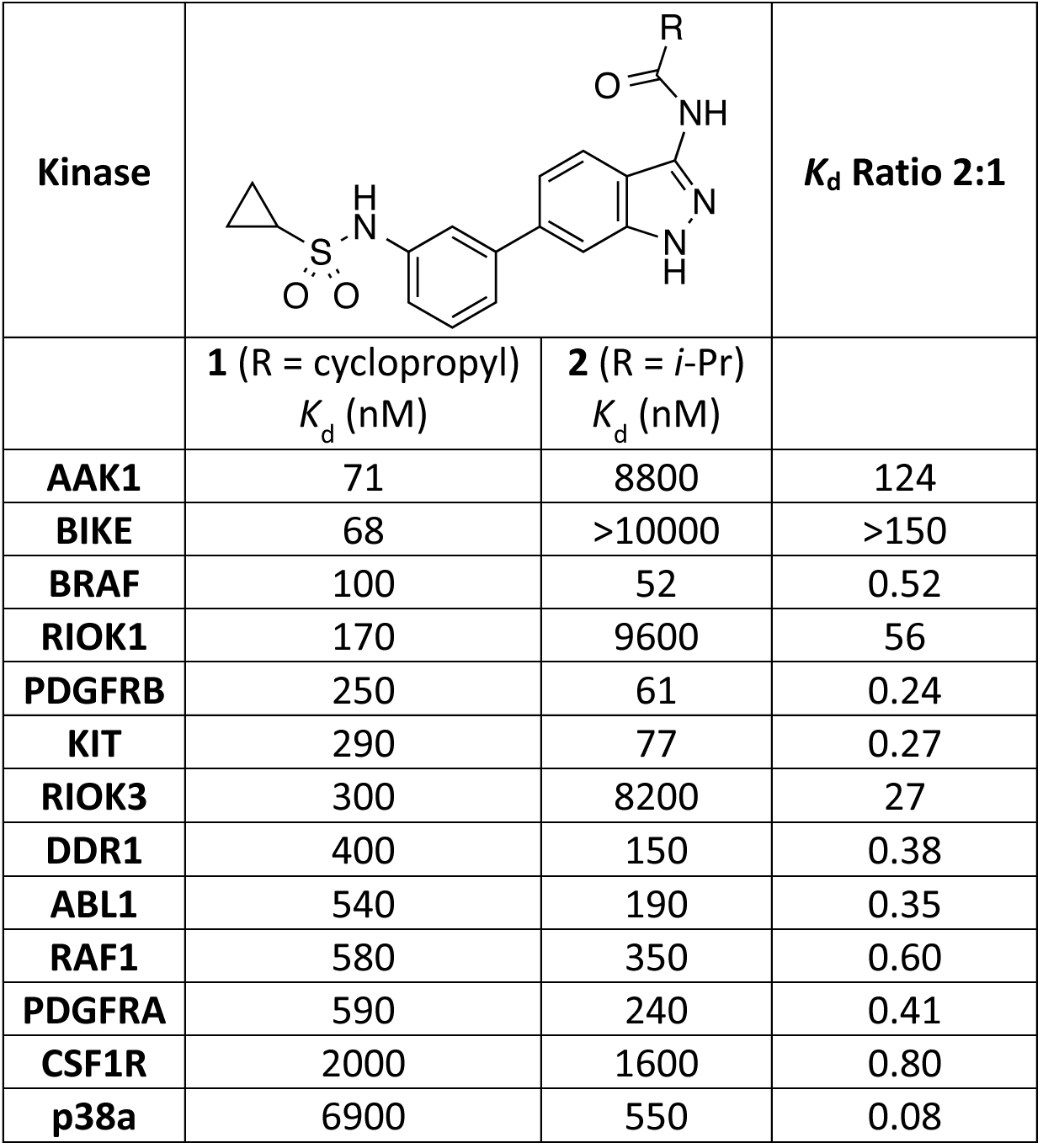
*K*_d_ values for selected kinases. *K*_d_ values were determined for kinases observed to have >70% displacement of immobilized ligand with either **1** or **2** in a KINOMEscan screen at 1 μM compound.

| | **1** (R = cyclopropyl)<br>$K_d$ (nM) | **2** (R = *i*-Pr)<br>$K_d$ (nM) | |
| --- | --- | --- | --- |
| **AAK1** | 71 | 8800 | 124 |
| **BIKE** | 68 | >10000 | >150 |
| **BRAF** | 100 | 52 | 0.52 |
| **RIOK1** | 170 | 9600 | 56 |
| **PDGFRB** | 250 | 61 | 0.24 |
| **KIT** | 290 | 77 | 0.27 |
| **RIOK3** | 300 | 8200 | 27 |
| **DDR1** | 400 | 150 | 0.38 |
| **ABL1** | 540 | 190 | 0.35 |
| **RAF1** | 580 | 350 | 0.60 |
| **PDGFRA** | 590 | 240 | 0.41 |
| **CSF1R** | 2000 | 1600 | 0.80 |
| **p38a** | 6900 | 550 | 0.08 |

AAK1 and BIKE, along with the related GAK and STK16, comprise the Numb-associated kinase (NAK) subfamily of protein kinases [8]. NAK family members have been proposed as potential drug targets for a wide range of therapeutic areas, including virology, oncology, neurology, and ophthalmology [9–12]. In contrast, relatively little is known about the relevance to cellular processes of the atypical kinases RIOK1 and RIOK3. Accordingly, to explore the molecular basis of the remarkable differential activities of **1** and **2**, we decided to study their interaction with the NAK family kinases. **1** and **2** were evaluated in FRET-based ligand binding displacement assays for all four NAK family members [13]. The cyclopropyl analog **1** showed >300-fold higher affinity for AAK1 and BIKE relative to the isopropyl analog **2** (**Table 2**). Both compounds demonstrated weaker affinity for GAK and STK16, although a >50-fold preference for **1** relative to **2** was observed with the latter kinase.

**Table 2.** NAK family FRET-based binding displacement assay results.

| Kinase | $K_i$ (nM) | | Ratio |
| --- | --- | --- | --- |
|  | 1 | 2 |  |
| AAK1 | 8.3 | 2700 | 320 |
| BIKE | 13 | 3100 | 240 |
| GAK | 1600 | 4900 | 3 |
| STK16 | 330 | 19000 | 58 |

### Structural analysis of compound 1 binding

BIKE and AAK1 are the closest members within the NAK family and share a high sequence identity—74% over their kinase domains and 81% within their ATP-binding sites. In order to obtain insights into the molecular details of ligand interactions with these kinases, we determined the cocrystal structure of BIKE kinase domain bound to compound **1** (**Table 3**), reasoning that both kinases are likely to interact with **1** and **2** in similar manners.

**Table 3.** Crystal structure data collection and refinement statistics.

|  |  |
| --- | --- |
| Protein | BIKE |
| Ligand | **1** |
| X-ray source | DLS I02 |
| a (Å) | 41.96 |
| b (Å) | 111.15 |
| c (Å) | 164.05 |
| $\alpha, \beta, \gamma$ (°) | 90, 90, 90 |
| Space group | I 2 2 2 |
| Wavelength (Å) | 0.97949 |
| Resolution limit (Å)* | 28.6-2.41 (2.50-2.41) |
| Number of observations* | 81.078 (7.855) |
| Completeness (%)* | 99.1 (95.2) |
| Multiplicity* | 5.3 (5.2) |
| Rmerge (%)* | 0.074 (0.49) |
| Mean I/σI * | 13.2 (3.1) |
| Refinement |  |
| Resolution Range (Å) | 21.75-2.41 |
| Rcryst | 0.176 |
| Rfree | 0.216 |
| No. protein atoms | 2,317 |
| No. ligand atoms | 28 |
| No. solvent atoms | 89 |
| Mean B-factor (Å) | 53.1 |
| Rmsd bond lengths (Å) | 0.010 |
| Rmsd bond angles (°) | 1.04 |
| Ramachandran statistics (%) |  |
| Favored | 98.3 |
| Allowed | 1.7 |
| Outlier | 0 |
| PDB ID | 5IKW |
| Crystallization conditions | 10% (v/v) Broad MW PEG smear,<br>3.2 M MgCl<sub>2</sub>, 100 mM Hepes pH<br>7.0 |

\* Highest resolution shell shown in parentheses.

* Highest resolution shell shown in parentheses.

BIKE-**1** crystals diffracted to 2.4 Å and contained a single protein chain per asymmetric unit (chain A). Sorrell and colleagues have recently reported the structural characterization of all four NAK family member kinase domains, including BIKE and AAK1 [8]. As previously observed, our structure of BIKE displayed a catalytically-competent conformation in which residues in the protein regulatory spine (“R spine”; residues M99, Y111, H178 and F199) are aligned and side-chain atoms from conserved residues in β3 and α-C (K79 and E95, respectively) form an ion pair [14, 15]. Moreover, the NAK family-exclusive helical domain (P209 to Y224) found C-terminal of the activation segment (from ^198^DFG to APE^233^ domains), dubbed ASCH (activation segment C-terminal helix) [8, 16, 17], also displayed an identical conformation in the BIKE structure described here (**Figure 1a**).

**Figure 1.**
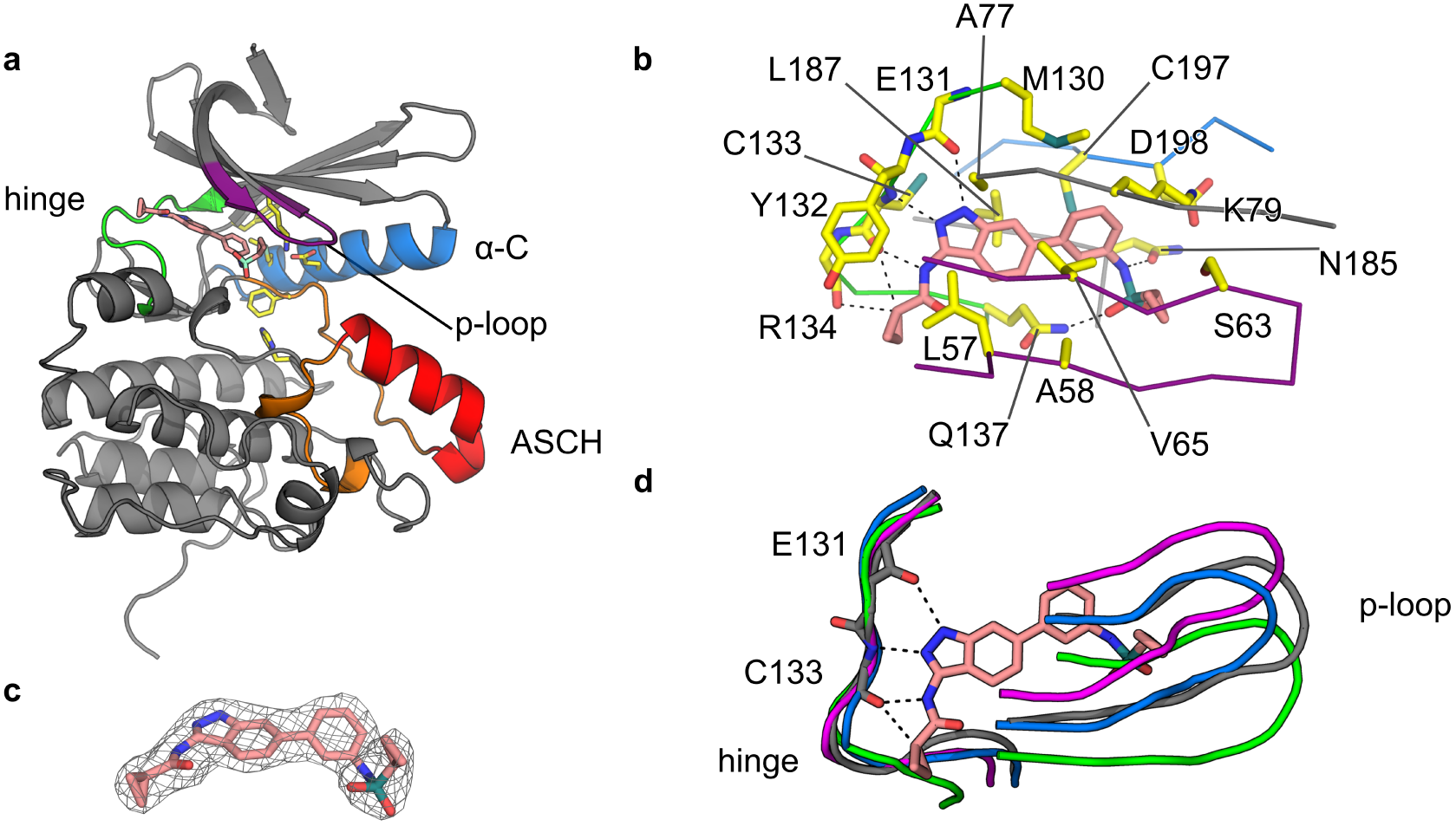
Crystallographic characterization of BIKE bound to 3-acylaminoindazole **1.** (**a**) BIKE-**1** adopts an active conformation. Protein (cartoon) regions important for activity and ligand interaction are indicated and highlighted in different colors. As for other NAK-family members, the activation segment of BIKE (orange) contains a C-terminal helical segment (ASCH; in red). Inhibitor **1** (carbon atoms in pink) and BIKE residues (carbon atoms in yellow) taking part in the regulatory hydrophobic spine and the critical ion pair leading to an active conformation are shown as sticks. (**b**) Details of the BIKE-**1** interactions. Inhibitor **1** is anchored to the hinge (green ribbon) via hydrogen bonds (dashed lines) and complements the hydrophobic cavity formed by aliphatic side chains from p-loop (purple ribbon) and β-6 residues. Potential interactions between main chain carbonyl and the cyclopropane group are indicated by dashed lines. Hydrogen bonds between Asn185, Gln137 and the sulfonamide group are also shown (dashed lines). (**c**) Superposition of NAK family proteins AAK1 (blue; PDB ID 4WSQ), BIKE (grey, PDB IDs 5IKW, this work), GAK (PDB ID 4Y8D) and MPSK1 (PDB ID 2BUJ) shows variation in the p-loop region despite good alignment in the hinge region. Compound anchoring hydrogen bonds are shown for BIKE and **1** (sticks). (**d**) Electron density (2Fo-Fc) maps (grey mash; 1.0 σ contour) for **1**. (**e**) Primary sequence comparison of NAK family hinge and P-loop regions.

The cocrystal structure of **1** and BIKE reveals that 3-acylaminoindazole **1** is anchored to the protein hinge region (BIKE residues 131-137) *via* hydrogen bonds between the N atoms of the indazole group and main chain carbonyl and NH groups of residues G131 and C133, respectively. An additional hydrogen bond is made between the amino group of **1** and the carbonyl group of C133. At the other end of the inhibitor, the sulfonamide group engages residues Q137 and N185 via hydrogen bonds. These interactions position the pendant aliphatic group into a hydrophobic cavity within the P-loop formed by residues V65 and S65. The indazole and the aniline rings are not coplanar to each other. This flexibility provides excellent shape-complementarity between the inhibitor and the ATP-binding pocket; especially with the aliphatic side chains from L187 and from p-loop residues L57, A58 and V65. There is also a sulfur-π interaction between the aniline ring and the sulfur atom of C197 (**Figure 1b**). Residue C197 is adjacent to the protein DFG motif and has been proposed as a site of covalent modification by the inhibitor (5*Z*)-7-oxozeaenol [8]. The cyclopropane group adjacent to the carboxyl group in **1** is accommodated within a sharp turn of the peptide backbone involving residues occupying positions +3, +4 and +5 from the so-called gatekeeper, or GK, residue (M130 in BIKE). This kink in the peptide backbone allows the carbonyl groups of residues GK+3 (C133) and GK+4 (R134) to point towards the cyclopropyl C-H group, approaching at a distance of 3.6 and 3.3 Å, respectively (**Figure 1b**).

We compared our BIKE-**1** structure with published structures of DDR1 (PDB: 5FDP), BRAF (PDB: 4XV9), AAK1 (PDB: 4WSQ), and RIOK1 (PDB: 4OTP) to attempt to rationalize the structural basis differential activity of **1** and **2** on some but not all kinases. Surface electrostatic potential is quite different for kinases that show differential activity compared with those that do not. AAK1, BIKE, and RIOK1 have bland surface charge where the critical cyclopropyl moiety of **1** is accommodated within the binding pocket. In contrast, BRAF and DDR1 have highly charged surfaces around their corresponding hinge binding regions.

Superimposing our BIKE-**1** structure with the previously published structures from the NAK family [8] by aligning the hinge residues (rmsd < 0.3 Å for 7 equivalent Cα) revealed that the P-loop of STK16 and GAK are substantially displaced compared to the position occupied by this region in BIKE and AAK1 structures (**Figure 1d**). The extensive contacts **1** makes with the P-loop may explain its selectivity for BIKE and AAK1 over the other two members of the NAK family. Interestingly, the primary sequence alignment of NAK family members reveals that sequence conservation is high for the P-loop region and low for the hinge moiety (**Figure 1e**).

### Computational studies

A range of computational methodologies was applied to shed light on the differential affinities of **1** and **2**. Energy minimization calculations in the unbound state using the OPLS3 force field [18] and the GB/SA [19] solvation model suggest that dehydration penalty upon binding is not a dominant factor, as similar hydration values were obtained for **1** (-29.8 kcal/mol) versus **2** (-29.2 kcal/mol); the cyclopropyl analog was actually slightly more penalized. To investigate the electronic properties, electrostatic potential derived atomic charges using the B3LYP exchange-correlation energy functional [20] and the 6-31G** basis set [21] were obtained in the gas phase. In this case, striking differences were observed for the C-H bond adjacent to the carbonyl group; it was much more polarized for cyclopropyl than isopropyl (**Figure 2**), with no major differences for the adjacent amide.

**Figure 2.**
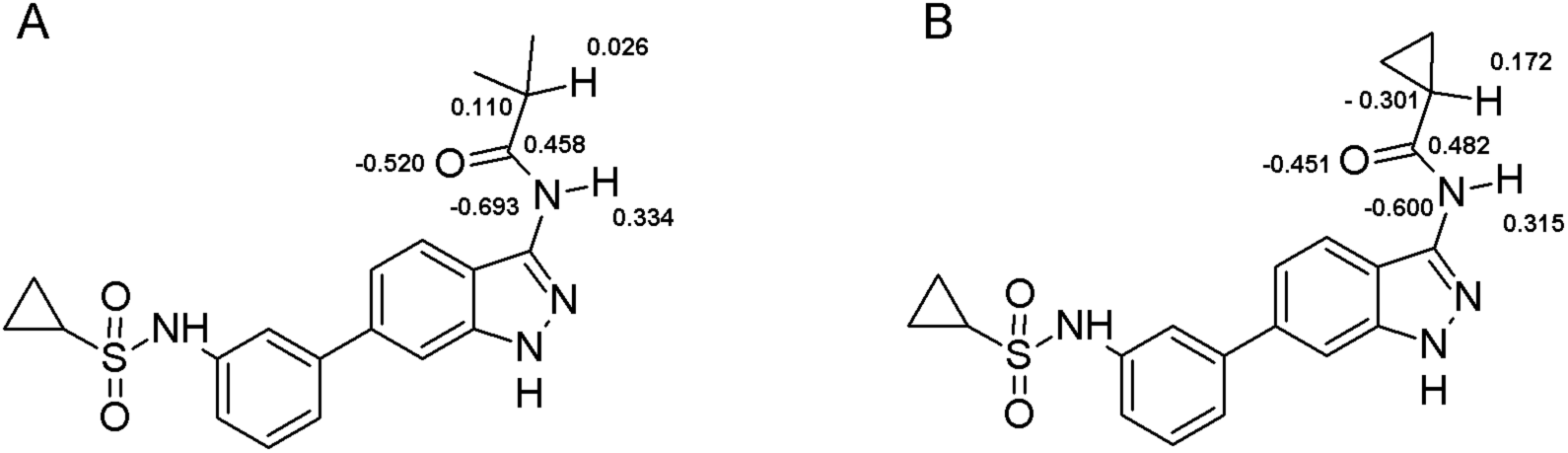
Computationally generated electrostatic potential (ESP) derived atomic charges for (A) **2** and (B) **1**.

Beyond defining potential enthalpic differences between cyclopropyl and isopropyl substituents, we also investigated possible entropic differences between **1** and **2** for selected kinases. The Glide docking application was used to predict binding poses in each investigated kinase, and the components of entropy change (translational, rotational, and vibrational) upon binding of **1** and **2** to the kinases were calculated using the RRHO approximation (**Table 4**). Translational and rotational entropic losses were essentially equivalent for **1** and **2** at any given kinase. However, the vibrational entropic gain was higher for **1** with all kinases except p38α. Interestingly, upon binding, the largest overall changes in entropy, or Δ (T ΔS_*bind*_), between **1** and **2** were observed for AAK**1** and BIKE. These two kinases displayed the two largest K_d_ ratios. Δ (TΔS_*bind*_) between **1** and **2** was markedly lower for those kinases not displaying significant activity cliffs. However, there are two exceptions to this observation, namely KIT and RIOK1. The former displayed a significant Δ(TΔS_*bind*_) between **1** and **2**, although there is not a significant difference in affinity. With RIOK**1**, the entropic difference between **1** and **2** was modest, despite a K_d_ ratio **2:1** of 56 (**Table 1**).

**Table 4.**
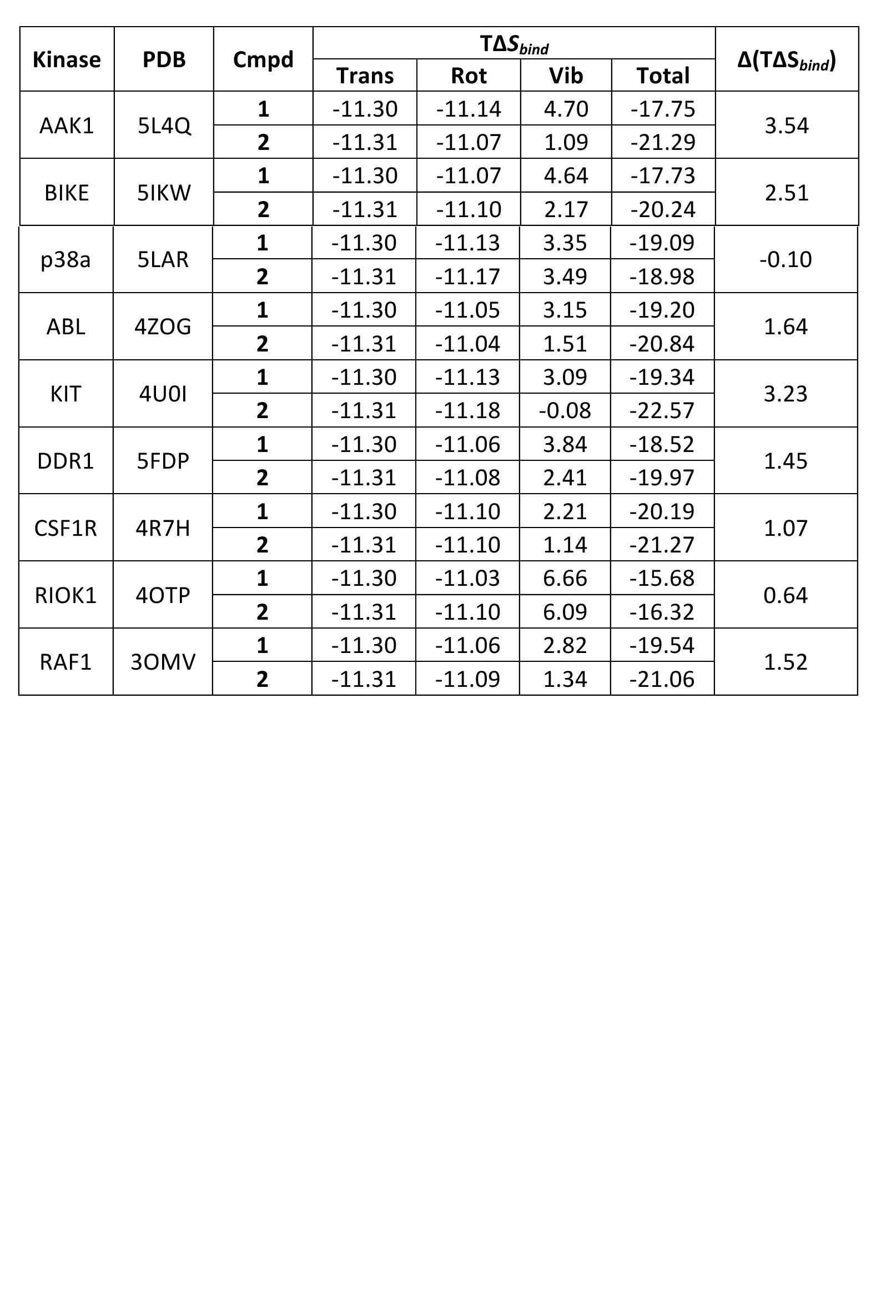
Thermodynamic parameters of binding **1** and **2** with kinases.

## DISCUSSION

The term “activity cliff” has been proposed to describe pairs of structurally similar compounds, like **1** and **2**, with a large difference in potency relative to a common target [22]. There have been other reports of kinase inhibitor activity cliffs involving cyclopropyl and isopropyl carboxamides proximal to their hinge binding moiety (**Table 5**). The 2-cyclopropylcarboxamidopyridine **3a** showed >100x higher affinity for TYK2 than the isopropyl analog **3b** [23]. This effect is not limited to binding assays as triazolopyrimidines **4** show >100x difference IC_50_ value in a JAK1 biochemical activity assay [24]. Similarly, **5a** displayed a much higher inhibition potency than **5b** in an enzymatic activity assay of mutant B-Raf^V600E^ activity [25]. These examples substantiate our observations that subtle changes in structure can lead to selective inhibition of specific protein kinases. To the best of our knowledge, **3-5** were not subjected to broad kinome activity characterization, and the breadth of the associated activity cliffs has not been evaluated.

**Table 5.**
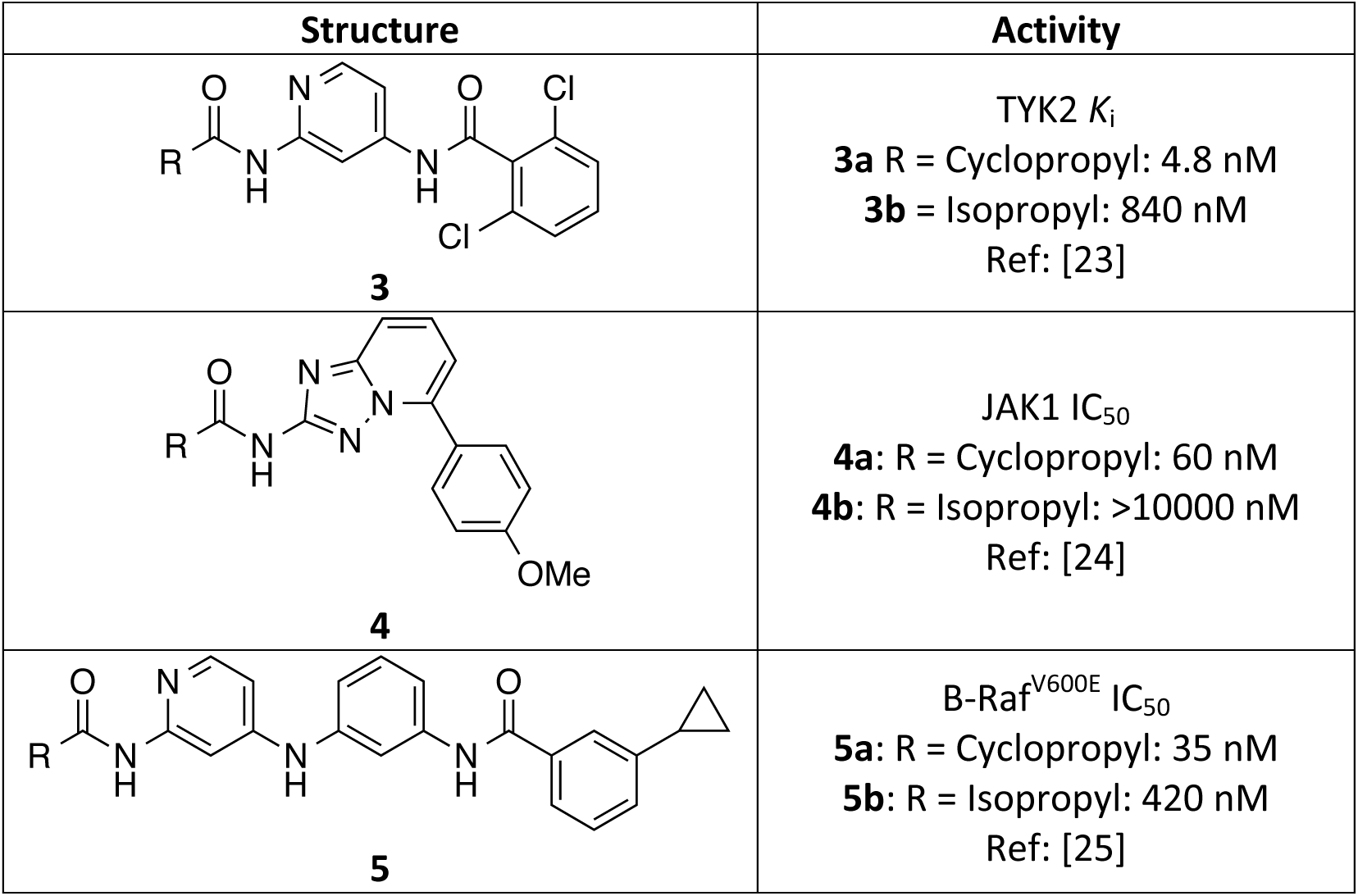
Previously reported matched pairs of kinase inhibitors demonstrating an affinity or activity cliff between cyclopropyl and isopropyl compounds.

| Structure | Activity |
| --- | --- |
| **3** | TYK2 $K_i$<br>**3a** R = Cyclopropyl: 4.8 nM<br>**3b** R = Isopropyl: 840 nM<br>Ref: [23] |
| **4** | JAK1 $IC_{50}$<br>**4a**: R = Cyclopropyl: 60 nM<br>**4b**: R = Isopropyl: >10000 nM<br>Ref: [24] |
| **5** | B-Raf<sup>V600E</sup> $IC_{50}$<br>**5a**: R = Cyclopropyl: 35 nM<br>**5b**: R = Isopropyl: 420 nM<br>Ref: [25] |

The cyclopropyl ring is a versatile lipophilic group that frequently appears in preclinical and clinical candidate molecules. It has been described to have “magical” properties that can enhance potency, reduce off-target effects, increase metabolic stability and brain permeability, yet the precise molecular basis for these effects remains poorly understood [26]. Theoretical studies on cyclopropane and its derivatives suggest that atomic bonding results mainly from the overlap of three *sp*^2^ hybrids (one on each carbon) pointing towards the center of the ring. Thus, the electron distribution in the C-C internuclear region is not concentrated along the line between the nuclei, but rather slightly outside this line. As a consequence, the C-C bonds have enhanced π character, and the C-H bonds have more *s* character. In other words, the cyclopropyl ring behaves much like an alkene, interacts with neighboring π-electron systems, and has CH bonds with increased acidity relative to standard alkanes [27]. These effects are apparent when the emerging negative charge on the C-H carbon is stabilized by conjugation, such as the one between the π-like cyclopropyl and an amide carbonyl group in **1**. Calculated electrostatic potential charges, as demonstrated above, may contribute to the activity difference between **1** and **2**.

3-Acylaminoindazoles are a known class of ATP-competitive protein kinase inhibitors [28]. The heterocyclic core mimics the adenosine base of ATP and forms hydrogen bonds with the hinge motif of the enzyme. **1** and **2**, differing by only two hydrogen atoms were screened across 400 human protein kinases to define both compounds as narrow spectrum inhibitors, with high affinity binding to <3% of the kinases surveyed. Both **1** and **2** bound to seven kinases (ABL1, BRAF, DDR1, KIT, PDGFRA, PDGFRB, and RAF1) with submicromolar affinity. Surprisingly, cyclopropyl amide **1** bound to an additional four kinases (AAK1, BIKE, RIOK1, and RIOK3) where isopropyl amide **2** was not effective. Thus, this pair of ATP-competitive kinase inhibitors have the potential to be used to probe selectively the biology of just 4 protein kinases while controlling for over 400 others in the human genome.

BIKE and AAK1 are both members of the NAK (Numb-associated kinase) family of protein kinases and share close sequence identity within their catalytic sites. Both kinases are potently inhibited by the 3-acylaminoindazole **1** and are likely to interact with the inhibitor through conserved interactions. The BIKE-**1** cocrystal structure provides a clear view into the molecular details of the enzyme-ligand interaction. The critical cyclopropyl carboxamide sits in a pocket adjacent to the hinge-binding residues and is orientated toward the protein surface. The amide NH of **1** and the cyclopropyl ring C-H sit in close proximity (2.70 and 3.29 Å, respectively) to the Cys133 carbonyl oxygen (**Figure 3**). Our quantum mechanical calculations showed that the cyclopropyl group has an increased polarization across the ring C-H compared the corresponding isopropyl analogue. The complex between the cyclopropyl derivative **1** and BIKE is indicative of a CH-O nonconventional hydrogen bond. Electrostatic interactions dominate hydrogen bonds and the increased polarization of the cyclopropyl C-H bond, as evidenced by the electrostatic potential charges, may explain why **1** shows significantly higher affinity for AAK1 and BIKE than the isopropyl analog **2** but leaves unanswered the question about why **1** and **2** are equipotent at other kinases.

**Figure 3.**
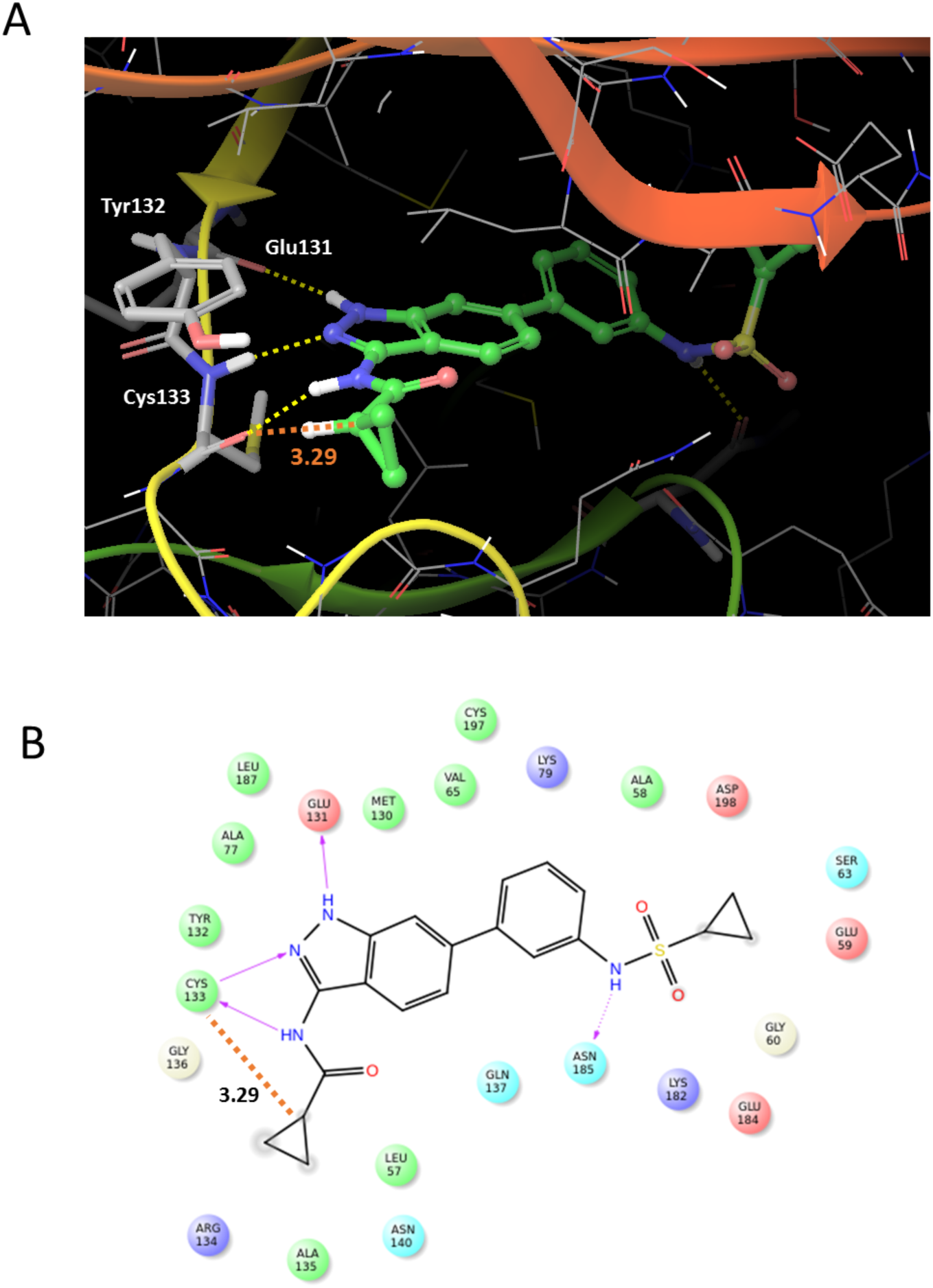
(A) **1** bound in BIKE active site. (B) Two dimensional compound interaction diagram of **1** and BIKE depicting adjacent residues and key interactions.

Modulating entropy contributions has previously been proposed as a means to increase binding affinity between small molecules and proteins [29]. Upon binding, ligands lose degrees of freedom with respect to translational, rotational, and vibrational motion.

This loss of freedom constitutes a loss of entropy, which disfavors binding. Given its triangular nature and smaller size, the cyclopropyl group may vibrate more freely within the protein, which favor binding when compared to the isopropyl group. Importantly, depending on the protein environment and architecture, this effect can be more pronounced. In our cocrystal structure of BIKE and **1**, the cyclopropyl group of interest is more solvent exposed. As such, this group is not as restricted by the binding site and does not experience a significant loss in vibrational entropy, which may be driving the activity cliff observed between **1** and **2.** Kinases for which an activity cliff between **1** and **2** is not observed may have the cyclopropyl group less solvent exposed, leading to a larger vibrational entropy loss.

We have suggested that differences in electrostatic potential or in vibrational entropy changes may help to rationalize the activity cliffs between **1** and **2** in some kinases but not in others. We recognize that there are myriad other subtle changes in the binding pockets that must also be explored to enable prediction of which kinases can be differentially inhibited by matched ligand pairs like **1** and **2**. Our enhanced understanding of the differential interactions of **1** and **2** within the binding pockets of AAK1, BIKE, and other kinases will improve our ability to produce narrow spectrum probes with optimal negative control analogs for use in biological studies.

## Supporting Information

**S1 Table. Results of KINOMEscan and select *K*_d_ determinations for 1 and 2**. KINOMEscan assay panel and associated *K*_d_ determinations were conducted at DiscoverX Corporation. Compounds were evaluated as single measurements at 1 μM. Displacement percentage (“Disp %”) reflects the extent to which the compound displaced the kinase construct from an immobilized ligand. *K*_d_ values were determined for all kinases with ≥70% displacement with **1** in the KINOMEscan screen at 1 μM compound and selected other kinases.

## MATERIALS AND METHODS

*N*-(6-(3-(cyclopropanesulfonamido)phenyl)-1*H*-indazol-3-yl)cyclopropanecarboxamide (**1**). ^1^H NMR (400 MHz, Methanol-*d*_4_) δ 7.83 – 7.78 (d, *J* = 8.6 Hz, 1H), 7.61 – 7.54 (m, 2H), 7.46 – 7.30 (m, 3H), 7.29 – 7.23 (ddd, *J* = 7.7, 2.2, 1.3 Hz, 1H), 2.62 – 2.49 (tt, *J* = 8.0, 4.8 Hz, 1H), 1.95 – 1.84 (ddd, *J* = 12.5, 7.9, 4.2 Hz, 1H), 1.07 – 0.77 (m, 8H). MS+ (– ES API) - 397.1

*N*-(6-(3-(cyclopropanesulfonamido)phenyl)-1*H*-indazol-3-yl)isobutyramide (**2**). ^1^H NMR (400 MHz, Methanol-*d*_4_) δ 7.82 – 7.75 (d, *J* = 8.6 Hz, 1H), 7.62 – 7.54 (dt, *J* = 10.9, 1.5 Hz, 2H), 7.47 – 7.31 (m, 3H), 7.31 – 7.22 (ddd, *J* = 7.8, 2.2, 1.3 Hz, 1H), 2.84 – 2.69 (hept, *J* = 6.8 Hz, 1H), 2.61 – 2.50 (tt, *J* = 8.0, 4.8 Hz, 1H), 1.29 – 1.15 (d, *J* = 6.8 Hz, 6H), 1.14 – 0.82 (m, 4H). MS+ (– ES API) - 399.1

### DiscoverX kinase affinity measurements

KINOMEscan assay panel profiling and associated *K*_d_ determinations were obtained at DiscoverX Corporation using their previously described methodology [30, 31].

### Binding displacement assays

Inhibitor binding was determined using a binding-displacement assay, which tests the ability of the inhibitors to displace a fluorescent tracer compound from the ATP binding site of the kinase domain. Inhibitors were dissolved in DMSO and dispensed as 16-point, 2x serial dilutions in duplicate into black multiwell plates (Greiner). Each well contained either 0.5 nM or 1 nM biotinylated kinase domain protein ligated to streptavidin-Tb-cryptate (Cisbio), 12.5 nM or 25 nM Kinase Tracer 236 (ThermoFisher Scientific), 10 mM Hepes pH 7.5, 150 mM NaCl, 2 mM DTT, 0.01% BSA, 0.01% Tween-20. Final assay volume for each data point was 5 μL, and final DMSO concentration was 1%. The plate was incubated at room temperature for 1.5 hours and then read using a TR-FRET protocol on a PheraStarFS plate reader (BMG Labtech). The data was normalized to 0% and 100% inhibition control values and fitted to a four parameter dose-response binding curve in GraphPad Software. The determined IC_50_ values were converted to *K*_i_ values using the Cheng-Prusoff equation and the concentration and *K_d_* values for the tracer (previously determined).

### Cloning, Expression, Purification and crystallization

BIKE_38-345 (K320A, K321A)_ with a tobacco etch virus (TEV) protease cleavable, N-terminal 6xHis tag was expressed from vector pNIC-ZB. To improve BIKE crystallization, a cluster of surface entropy reduction mutations [32] was engineered into the expression construct, K320A/K321A [8]. For protein production, the host *E. coli* strain, BL21(DE3)-R3 expressing lambda phosphatase, was cultivated in TB medium (+ 50 μg/ml kanamycin, 35 μg/ml chloramphenicol) at 37°C until OD_600_ reached ^~^3 and then cooled to 18°C for 1 hour. Isopropyl 1-thio-D-galactopyranoside was added to 0.1 mM, and growth continued at 18°C overnight. The cells were collected by centrifugation then resuspended in 2x lysis buffer (100 mM HEPES buffer, pH 7.5, 1.0 M NaCl, 20 mM imidazole, 1.0 mM tris(2- carboxyethyl)phosphine, 2x Protease Inhibitors Cocktail Set VII (Calbiochem, 1/1000 dilution) and flash-frozen in liquid nitrogen. Cells were lysed by sonication on ice. The resulting proteins were purified using Ni-Sepharose resin (GE Healthcare) and eluted stepwise in binding buffer with 300 mM imidazole. Removal of the hexahistidine tag was performed at 4°C overnight using recombinant TEV protease. Proteins were further purified using reverse affinity chromatography on Ni-Sepharose followed by gel filtration (Superdex 200 16/60, GE Healthcare). Protein in gel filtration buffer (25 mM HEPES, 500 mM NaCl, 0.5 mM TCEP, 5% [v/v] glycerol) was concentrated to 12 mg/ml using 30 kDa MWCO centrifugal concentrators (Millipore) at 4°C. 3-acylaminoindazole **1** in 100% DMSO was added to the protein in a 3-fold molar excess and incubated on ice for approximately 30 minutes. The mixture was centrifuged at 14,000 rpm for 10 minutes at 4°C prior to setting up 150 nL volume sitting drops at three ratios (2:1, 1:1, or 1:2 protein-inhibitor complex to reservoir solution). Crystallization experiments were performed at 20°C. Crystals were cryoprotected in mother liquor supplemented with 20–25% glycerol before flash-freezing in liquid nitrogen for data collection. Diffraction data were collected at the Diamond Light Source. The best diffracting crystals grew under the conditions described in **Table 1.** When noted, crystal optimization used Newman’s buffer system [33] and defined PEG smears [34].

### Structure Solution and Refinement

Diffraction data were integrated using XDS [35] and scaled using AIMLESS from the CCP4 software suite [36]. Molecular replacement for was performed with Phaser [37] using BIKE/AZD7762 (PDB ID: 4W9W) [8] as a search model. Automated model building was performed with Buccanner [38]. Automated refinement was performed in PHENIX [39]. Coot [40] was used for manual model building and refinement, Structure validation was performed using MolProbity [41]. Structure factors and coordinates have been deposited in the PDB (see Table 1).

### Computation

Solvation free energies and quantum mechanical calculations were run using Macromodel ([42]) and Jaguar ([43]) from Schrodinger Inc. Further computational studies were performed using Maestro 11 from Schrodinger (Schrödinger Release 2016-4: Maestro, Schrödinger, LLC, New York, NY, 2016). The lowest energy configurations for compounds **1** and **2** at pH 7 were generated with the Ligand Preparation application (Schrödinger Release 2016-4: LigPrep, Schrödinger, LLC, New York, NY, 2016). Next, PDB crystal structures for AAK1, BIKE and the other kinases in Table 4 were downloaded and prepared for docking. Briefly, bond orders were assigned, hydrogen atoms were added, selenomethionine residues were converted to methionines, and missing side chains were filled with the Prime application. The protein grid for docking was set by selecting the center of the cocrystal ligand in the PDB structure. The Glide application was used for docking of 1 and 2 to all kinases (Schrödinger Release 2016-4: Glide, Schrödinger, LLC, New York, NY, 2016.). Flexible ligand sampling was allowed with the XP (extra precision) scoring function (http://pubs.acs.org/doi/abs/10.1021/jm051256o). Once the ligands were docked and the lowest energy poses were retrieved, a Rigid Rotor Harmonic Oscillation (RRHO) approximation was performed on the theoretical protein-ligand complex (https://www.ncbi.nlm.nih.gov/pmc/articles/PMC3329805/). For the RRHO approximation, an OPLS_2005 force field was applied (http://pubs.acs.org/doi/abs/10.1021/acs.jctc.5b00864).

## ACKNOWLEDGMENT

Aled Edwards is acknowledged for illuminating discussions and encouragement in the preparation of this manuscript. We thank Diamond Light Source for access to beamline I03 (proposal number MX14664) that contributed to the results presented here. The SGC is a registered charity (number 1097737) that receives funds from AbbVie, Bayer Pharma AG, Boehringer Ingelheim, Canada Foundation for Innovation, Eshelman Institute for Innovation, Genome Canada, Innovative Medicines Initiative (EU/EFPIA), Janssen, Merck & Co., Novartis Pharma AG, Ontario Ministry of Economic Development and Innovation, Pfizer, São Paulo Research Foundation-FAPESP, Takeda, and Wellcome Trust.

